# tTEscanR: A user-friendly integrative R package for quantifying and visualizing translation efficiency from sequencing data in diverse biological systems

**DOI:** 10.64898/2026.09.08.750101

**Authors:** Ana Varas-Sánchez, Carlos J. Gallardo-Dodd, Qun Li, William Gao, Markus Ringnér, Claudia Kutter

## Abstract

Graphical abstract

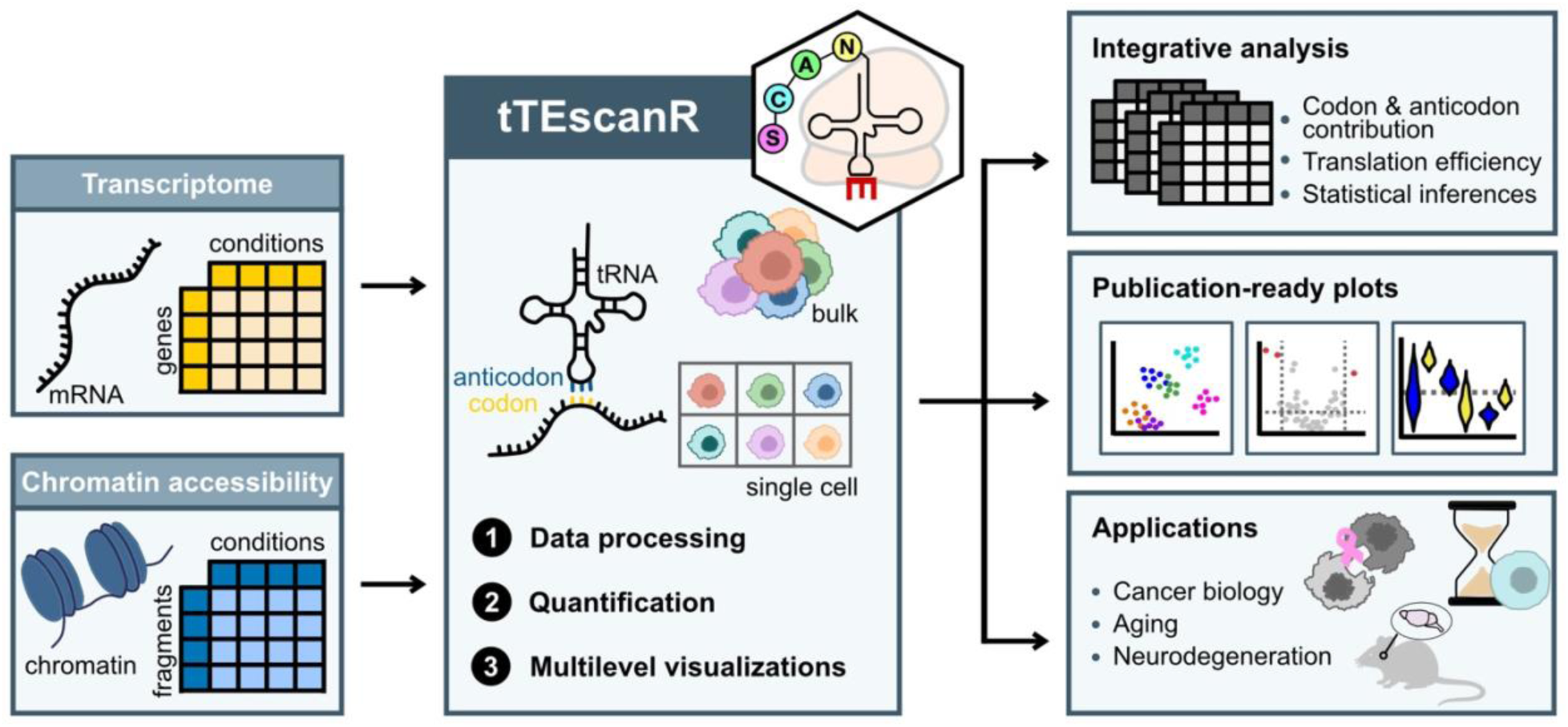

Translation elongation relies on accurate codon-anticodon pairing. Here, we present tTEscanR, an R package designed to investigate this translational interface. By quantifying mRNA codon demand alongside tRNA anticodon availability, tTEscanR provides scalable estimates of translation rates directly from standard transcriptomic and chromatin accessibility count matrices, where genomic features are represented as rows and experimental conditions as columns. tTEscanR seamlessly integrates into existing bulk and single-cell pipelines and provides modular functions for quality control, filtering, normalization, statistical analysis, and visualization. A structured data object centralizes workflow outputs and associated metadata. A built-in multilevel visualization module generates customizable, publication-ready graphical outputs to facilitate data interpretation and reproducibility of the complex translational landscape. We demonstrate the utility of tTEscanR across cancer biology, aging, and neurodegeneration datasets, uncovering critical translational regulatory programs overlooked by conventional analyses. tTEscanR is available in an open-source repository as a standalone tool or workflow plug-in.

## Background

The translation of messenger RNAs (mRNAs) into protein forms the critical link between the transcriptome and the proteome. Essential during this process are transfer RNAs (tRNAs) that act as adapters of the genetic code by pairing their anticodons with codons in mRNAs, thereby determining the precise amino acid sequence of proteins. Efficient decoding relies on a dynamic balance between the availability of anticodons in tRNA pools (supply) and the usage of corresponding codons in mRNA pools (demand), which together maintain translational homeostasis (1–4). When this balance is preserved, translation proceeds with high efficiency and fidelity; conversely, when disrupted, translational speed and accuracy may be compromised. Although this balance is clearly important, studying it systematically remains challenging.

A major obstacle lies in the need to accurately quantify both sides of this connection. In humans and mice, this involves relating approximately 20,000 protein-coding transcripts to a compact and redundant repertoire of 400 high-confidence tRNA genes (5). Importantly, tRNA genes are differentially active at any given time across cellular states, and multi-copy tRNA genes can exhibit highly unequal levels of expression (6–8).

To map the demand side, global codon usage is typically inferred from transcriptome-wide data by parsing coding sequences into triplets, computing gene-level frequencies, and weighting them by transcript abundance to reflect the actual translation pool. However, estimating the supply side of tRNA anticodon availably is considerably more complex (9). Mature tRNAs are not directly captured by standard RNA-seq because of their short length, extensive base modifications, stable secondary structure, and inefficient reverse transcription, resulting in substantial cross-platform variability among specialized tRNA profiling platforms (6,7,10–15). To overcome this limitation, several indirect strategies have instead leveraged Pol III occupancy (3,8,16–18) or chromatin accessibility as proxies for tRNA gene activity (19). Rather than measuring mature tRNA abundance, these approaches quantify tRNA gene usage, which can be aggregated at the tRNA anticodon and amino acid levels to estimate translational supply (17,20). Although these strategies for measuring codon demand and anticodon availability have become increasingly used, their implementations remain fragmented across custom scripts with limited modularity, standardization and interoperability. Consequently, this limits reproducibility and prevents systematic benchmarking across datasets, particularly in large-scale single-cell studies where sparsity and heterogeneity further complicate analysis.

While computational proxies offer a window into translational potential, the experimental gold standard for capturing active translation is ribosome profiling (Ribo-seq). By sequencing ribosome-protected mRNA fragments, Ribo-seq physically maps translating ribosomes across the transcriptome (21,22). Despite its high resolution, Ribo-seq requires high input material from bulk tissue and remains challenging to extend to single-cell applications (23,24). Thus, there is a growing need for scalable computational frameworks that can extract information about translational efficiency across the rapidly expanding universe of conventional transcriptomics and chromatin-accessibility datasets, including single-cell datasets for which dedicated measurements of translation are unavailable.

To overcome these experimental limitations and leverage widely available data, we previously established a computational concept that demonstrated how codon demand and tRNA anticodon supply can be jointly inferred from sequencing data and combined to estimate theoretical translation efficiency (tTE) (19). In this approach, chromatin accessibility at tRNA genes measured by ATAC-seq or single-cell ATAC-seq serves as a measure for tRNA expression and shows strong concordance with Pol III occupancy. By correlating transcript-level codon demand to the corresponding anticodon supply, we introduced tTE as a quantitative measure of the predicted compatibility between the mRNA codon pool and the available tRNA repertoire. Higher tTE scores indicate a more optimal balance between tRNA anticodon supply and mRNA codon demand. This work established the conceptual and computational basis for estimating translational constraints without directly measuring ribosome occupancy.

Here, we introduce tTEscanR (**t**heoretical **T**ranslation **E**fficiency **scan**ner implemented in **R**), an open-source R/Bioconductor package that transforms this previously established concept into a generalizable, standardized, and scalable framework for translational analysis. Rather than providing another implementation of tTE, tTEscanR addresses the broader methodological challenge of making codon-anticodon analysis reproducible and applicable across heterogeneous datasets. It brings the complete analysis into a unified framework, from (i) quantification and quality control of codon and tRNA gene usage through normalization, (ii) integration of transcriptomic and chromatin accessibility data to jointly characterize mRNA codon demand and tRNA gene activity within the same analytical framework, (iii) tTE calculation, (iv) statistical comparison, and (v) visualization. Importantly, tTEscanR introduces dedicated strategies for the sparse and heterogeneous measurements encountered in single-cell data, enabling robust analysis across individual cell populations as well as pseudobulk and population-level datasets. Thus, tTEscanR moves beyond a proof-of-concept estimation of tTE to provide a reproducible analytical infrastructure for systematically interrogating the translational landscape encoded in standard sequencing datasets.

We demonstrate the utility of tTEscanR across diverse datasets to characterize translational efficiency in cancer, neurodegeneration, and aging at single-cell resolution. These applications demonstrate that the framework can resolve differences in codon demand, tRNA gene activity, anticodon supply, and tTE across cell types and disease states. Our analyses uncovered previously unrecognized patterns of codon-anticodon adaptation, including cell-type-specific shifts in translational balance in pathological states, highlighting the dynamic nature of translation efficiency in complex biological systems. By providing a reproducible and scalable framework that integrates seamlessly with standard RNA-seq and ATAC-seq workflows, tTEscanR facilitated systematic investigation of translational efficiency and enabled the reuse of publicly available sequencing datasets to uncover novel translational mechanisms.

## Results

### tTEscanR enables accurate analysis of translation efficiency from sequencing data

tTEscanR is a powerful open-source R package available on Bioconductor for the standardized analysis of translation efficiency from bulk and single-cell sequencing data. It implements a computational framework that estimates tTE by integrating mRNA codon usage with tRNA anticodon availability. The package supports analyses across three interconnected layers of information: mRNA and tRNA gene usage, codon and anticodon pools, and amino acid demand and supply. This multi-layered approach provides a comprehensive view of translational dynamics across various biological contexts.

tTEscanR was designed to transform our previously developed methodology (19) into a comprehensive software platform. The R package addresses key limitations of our original scripts, and provides a standardized workflow, supporting modular analyses, replacing manually adapted scripts with scalable and reproducible analyses that integrate seamlessly with existing bioinformatic pipelines. tTEscanR supports both end-to-end workflows or independent execution of individual analytical components. This design facilitates integration with standard transcriptomic and epigenomic analysis pipelines while accommodating diverse data modalities and experimental designs. To maximize broad applicability, tTEscanR is fully compatible with both model and non-model organisms and supports custom genomic references.

The core tTEscanR workflow consists of six major analytical steps (**Figure 1**): (i) data acquisition and preprocessing; (ii) construction of a standardized tTEscanR object; (iii) quantification and analysis of codon and anticodon pools; (iv) estimation of amino acid demand and supply; (v) computation of the tTE scores; and (vi) downstream data interpretation through integrated statistical and visualization modules (**Methods**). To demonstrate the analytical scope and versatility of tTEscanR, we applied the package to two independent high-resolution multi-omic datasets representing distinct biological and experimental contexts. These analyses illustrate the applicability of tTEscanR across heterogenous biological systems and sequencing modalities.

**Figure 1.**
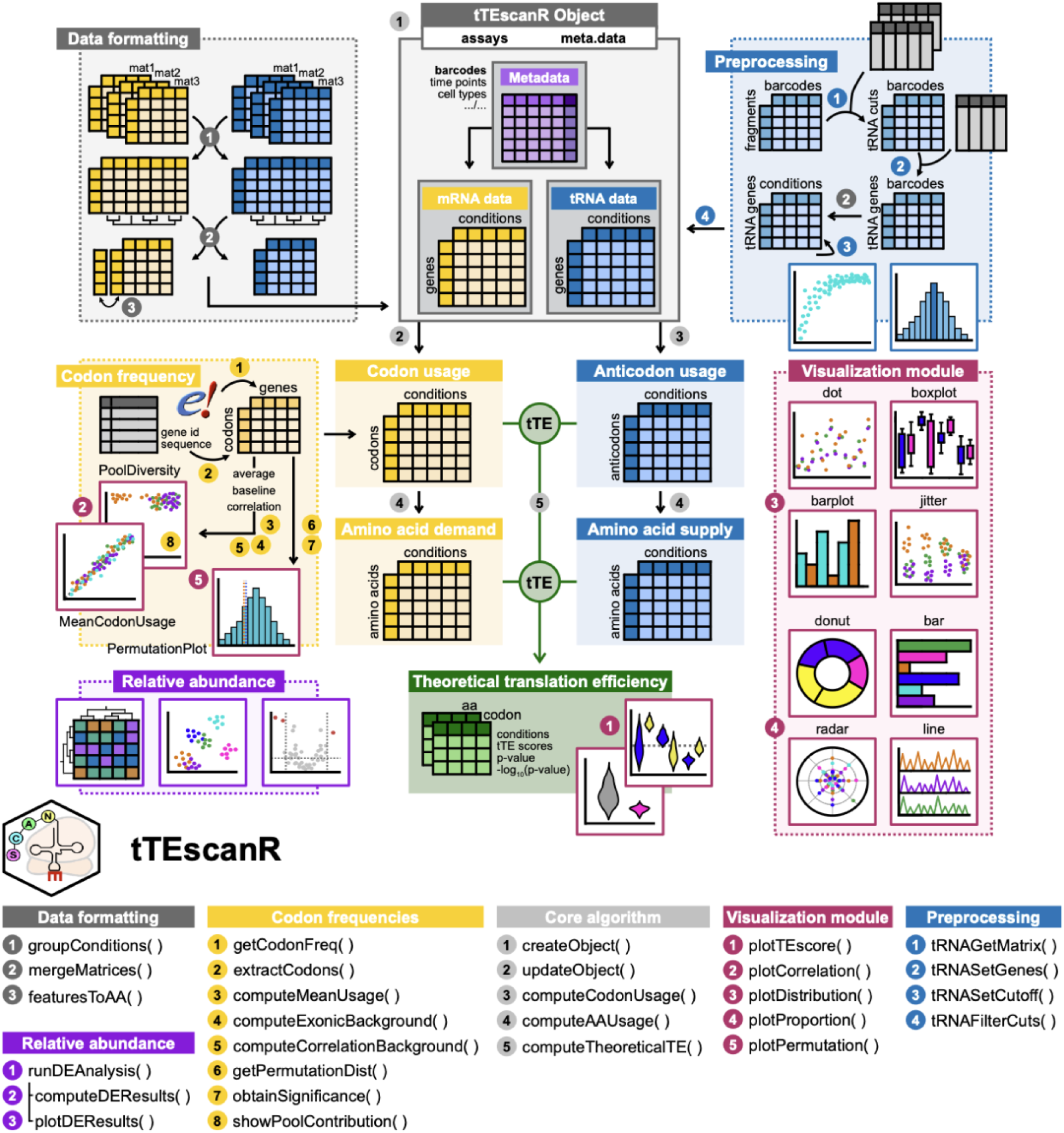
Schematic overview of the tTEscanR analytical pipeline. The tTEscanR pipeline consists of six major analytical steps: (i) data acquisition and preprocessing (top corner boxes); (ii) construction of a standardized tTEscanR object (top central box); (iii) quantification and analysis of codon (yellow-colored boxes) and anticodon pools (blue-colored boxes); (iv) estimation of amino acid demand (yellow-colored boxes) and supply (blue-colored boxes); (v) computation of tTE scores (green-colored box); and (vi) downstream data interpretation through integrated statistical (purple-colored box) and visualization modules (magenta-colored box). General plotting functions are grouped within the visualization module, whereas step-specific plots are integrated into their respective functional blocks. The tTEscanR object is structured into *assays* and *meta.data* sections, accommodating various feature count matrices as input. tTE scores can be computed across matching conditions in the mRNA (yellow) and tRNA (blue) datasets at either the codon-anticodon or amino acid demand-supply levels. Core algorithmic functions are housed within the central panel. Arrows indicate the direction of data flow through the pipeline, and primary user functions are listed below.

### Erythrocytes deviate from the otherwise stable codon and amino acid pools

We utilized a patient-derived pediatric acute myeloid leukemia (pAML) cohort (25) to evaluate translation dynamics across clinical disease stages. Following rigorous preprocessing and quality assessment (**Methods**), we retained a final cohort of 11 patients with pAML with matching diagnosis, remission, and relapse data **(Figure 2A; Supp. Figure 1)**. At the gene expression level, we were able to clearly separate normal (remission) and malignant (diagnosis and relapse) states, reflecting the transcriptional differences between leukemic and normal hematopoietic populations **(Figure 2B, left panel)**. Despite these pronounced gene expression changes across states, codon usage and amino acid demand remained remarkably stable across most cell populations, supporting previous observations that codon usage is largely conserved across species, development stages, tissues and cell types (1,6,8,19). However, dimensionality reduction analysis performed with tTEscanR identified erythrocytes as a notable exception to this overall stable pattern, as these cells formed a distinct cluster based on codon usage and amino acid demand that separated from the remaining hematopoietic populations **(Figure 2B, center and right panels)**.

**Figure 2.**
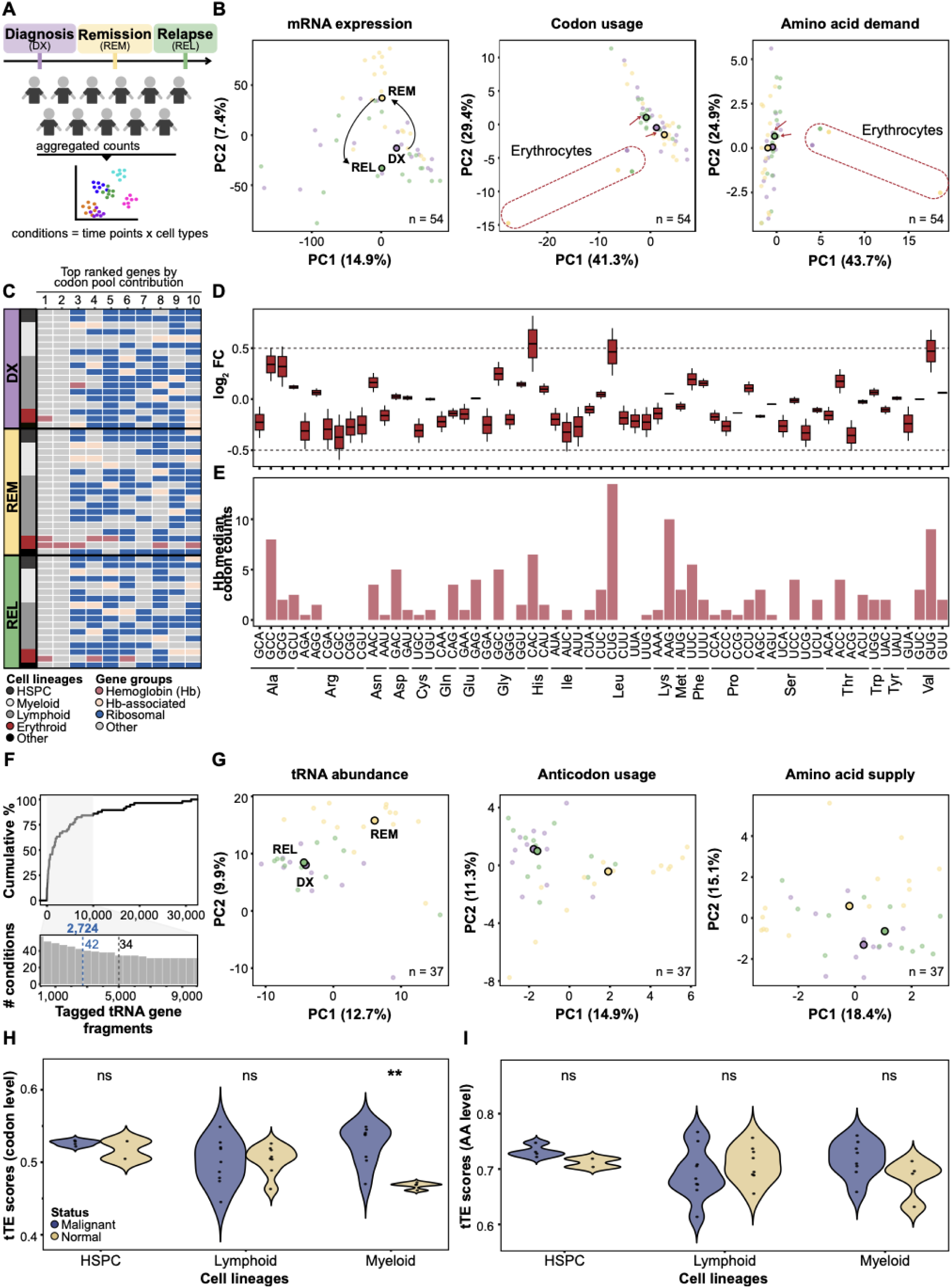
tTEscanR reveals cell lineage-specific translational profiles in pAML across clinical stages. **A.** Overview of the pediatric acute myeloid leukemia (pAML) cohort analyzed in this study. Paired scRNA-seq and scATAC-seq data were retrieved across three clinical stages: diagnosis (DX, purple), remission (REM, yellow) and relapse (REL, green). Individual cells of each patient are aggregated into cell types following a pseudobulk strategy. We profiled 19 cell types at each of the three clinical stages, yielding a total of 57 conditions before filtering out low-quality samples. **B.** PCA plots visualize mRNA gene expression (left), codon usage (center) and amino acid demand (right), colored by clinical stage. Each data point represents a pseudobulk condition defined by the combination of clinical stages and cell types (n = 54, after filtering). Median values are highlighted with larger circles. Red dashed outlines mark the erythroid cells, illustrating the deviation from the otherwise stable codon and amino acid demand pool. The red arrows mark erythroid cells that are not part of the dominant source of variation. **C.** Heatmap analysis of codon pool contribution across clinical stages and cell lineages. Each row corresponds to a specific condition (matching the individual data points in the PCA plots from panel B), showcasing the top 10 ranked genes contributing most heavily to codon usage. Erythrocytes (dark red) consistently feature hemoglobin genes (light red) in the top ranks. Highly abundant ribosomal genes (dark blue) are highlighted for comparison. **D.** Boxplot shows codon usage comparison between erythroid lineage (early and late erythrocytes) compared to all other cell types at the remission stage. Relative enrichment is plotted on the y-axis as log_2_ fold-change (FC). Statistical significance was defined using a threshold of log_2_ FC > 0.5. Three codons were significantly enriched in the erythroid lineage: Histidine-CAC, Leucine-CUG, and Valine-GUG. **E.** Barplot illustrates median codon counts across the ten hemoglobin complex genes present in the dataset (*HBA1*, *HBA2*, *HBB*, *HBD*, HBE1, *HBG1*, *HBG2*, *HBM*, *HBQ1* and *HBZ*). **F.** Data-driven selection of the optimal tRNA gene-associated threshold for the pAML dataset. The upper panel displays the cumulative percentage of retained conditions (y-axis) as a function of the number of tRNA gene-associated tagged fragments (x-axis) per condition. The lower panel magnifies the grey shaded region (1,000 to 9,000 tags) from the upper panel, with the y-axis denoting the actual number of conditions included. When multiple thresholds yield the same number of conditions, tTEscanR automatically selects the lowest threshold. Dashed lines indicate the dataset-specific threshold determined by tTEscanR (2,724 tags, light blue) and the previously recommended generic threshold (5,000 tags, grey). The number of retained conditions at each threshold (42 and 34, respectively) is indicated. **G.** PCA plots visualize tRNA gene abundance (left), anticodon usage (center) and amino acid supply (right), colored by clinical stage. Data was filtered using the dataset-specific tTEscanR threshold of 2,724 tRNA gene-associated tags determined in **(F)**. Erythroid cells were subsequently excluded, resulting in a total of 37 conditions for downstream analysis. Across all three levels of analysis (tRNA gene, anticodon, amino acid supply), malignant cells from the diagnosis (purple) and relapse (green) stages separated from normal hematopoietic populations in remission (yellow). **H-I.** Violin plots illustrate theoretical translation efficiency (tTE) scores across hematopoietic stem and progenitor (HSPC), myeloid, and lymphoid cell lineages. Normal cells from the remission stage (gold) are compared with malignant cells from the diagnosis and relapse stages (blue) at the **(H)** codon-anticodon and **(I)** amino acid demand-supply level. Asterisks indicate statistical significance (**, *p*-value < 0.01) derived from a two-tailed standard normal distribution (Z-test); ns, non-significant.

To identify the molecular basis of this erythrocyte-specific divergence, we used tTEscanR to quantify the contribution of individual transcripts to codon usage pattern (**Methods**). This analysis revealed that erythroid cells markedly reduced codon pool complexity by concentrating translational demand within a small number of highly expressed transcripts. In particular, hemoglobin genes dominated the erythroid transcriptome and imposed a strong demand for a specific subset of codons, thereby reshaping the global codon usage profile of these cells **(Figure 2C)**. Thus, rather than reflecting a generalized shift in translation, the erythroid codon landscape emerged directly from extreme transcriptional specialization. This observation closely mirrors the transcriptome-driven codon specialization previously reported in smooth muscle cells (19), where a limited number of highly expressed genes similarly drive codon usage. To further support this interpretation, we repeated the analysis after excluding erythrocytes. We found that removing this cell population largely abolished the variability observed across codon usage and amino acid demand profiles and restored the otherwise stable translational landscape shared by the remaining cell types **(Supp. Figure 2A)**.

While all stages contained a broad spectrum of hematopoietic cell types, remission samples lacked the overrepresentation of specific cell types that was observed in both diagnosis and relapse. Consequently, we focused on the remission stage, which contained a more balanced diversity of hematopoietic cell populations **(Supp. Figure 1C, top panel)**. Using tTEscanR, we quantified the distribution of individual codon frequencies across remission cell types and analyzed codon pools by comparing erythroid cells against all other cell lineages ( **Methods**). Consistent with the divergence observed in our dimensionality reduction analysis **(Figure 2B)**, cells of the erythroid lineage, spanning early to late maturation stages, displayed significant deviations in the usage of a restricted subset of codons **(Figure 2D; Supp. Figure 2B-C)**. Across multiple maturation stages, His-CAC, Leu-CUG, and Val-GUG codons were overrepresented in erythrocytes. To determine whether these shifts originated directly from the dominant hemoglobin gene transcripts **(Figure 2C)**, we computed the codon usage profiles for the genes within the hemoglobin complex present in our data (**Methods**). The resulting profiles closely matched the high-frequency codons enriched within the hemoglobin sequences **(Figure 2E)**, demonstrating that lineage-specific transcriptional programs can actively reshape codon demand.

### A dataset-specific tRNA cuts threshold enhances robustness of anticodon quantification

Reliable estimation of tRNA anticodon availability from scATAC-seq data remains challenging because chromatin accessibility signals at tRNA loci are inherently sparse. In our previous work, we established that a minimum of 5,000 aggregated tRNA gene-associated scATAC-seq cuts per condition provides stable anticodon usage estimates (19). However, applying a fixed threshold across datasets can unnecessarily reduce sample retention and limit downstream analysis, particularly in heterogenous and sparse single-cell data.

To address this limitation, we developed a dedicated threshold optimization module implemented in tTEscanR that identifies more flexible, dataset-specific cutoffs balancing signal robustness and data retention (**Methods**). The module systematically evaluates the stability of anticodon quantification across increasing tRNA gene cut thresholds and determines the optimal point at which measurements become reproducible while maximizing the number of retained conditions. This functionality provides a standardized and data-driven strategy for quality control of tRNA-derived measurements.

Applying this module to the pAML scATAC-seq dataset identified an optimal stability threshold of 2,724 tRNA gene-associated cuts across 392 tRNA genes **(Figure 2F; Supp. Figure 2D)**. Using this optimized threshold, tTEscanR retained 42 conditions (defined by the combination of clinical stages and cell types) for downstream analysis **(Supp. Figure 1C, bottom panel)**. In contrast, applying the previously recommended generic threshold of 5,000 cuts would have retained only 34 conditions. These results demonstrate that adaptive thresholding implemented in tTEscanR increases statistical power while preserving robust anticodon quantification.

Following this initial filtering, we excluded erythroid cells from the subsequent analyses. This population is highly heterogeneous, spanning multiple erythrocyte differentiation stages. As these cells undergo maturation and enucleation, they lose accessible nuclear chromatin, rendering scATAC-seq profiles unreliable. Additionally, this exclusion aligns with the filtering criteria applied to the scRNA-seq dataset, maintaining a matched cell type composition between both modalities. As a result, the final dataset comprised 37 matched conditions. Dimensionality reduction based on tRNA gene usage, anticodon, and amino acid supply separated malignant cells from diagnosis and relapse stages from normal hematopoietic populations in remission **(Figure 2G, Supp. Figure 2E)**, suggesting coordinated remodeling of the translational supply landscape during leukemic progression.

### Myeloid cells from pAML patients exhibit increased theoretical translation efficiency

We next used tTEscanR to determine whether the disease-associated differences translated into altered tTE across disease states. To this end, we quantified tTE scores across the 37 matched conditions shared between the scRNA-seq and scATAC-seq datasets **(Supp. Table 2)**. To identify pAML-associated translational signatures, we grouped malignant stages (diagnosis and relapse) and compared them against the remission-derived normal state. Our analysis revealed a statistically significant increase of tTE scores in malignant myeloid cells at the codon-anticodon level compared with remission cells **(Figure 2H)**, indicating a stronger coordination between codon demand and anticodon supply. Although our analysis at the amino acid level showed similar trends **(Figure 2I)**, it did not reach statistical significance, suggesting that the disease-associated signal emerges primarily at the codon-anticodon level, which more directly reflects the decoding interactions occurring at the ribosome during translation elongation. Collectively, these analyses demonstrate that tTEscanR sensitively captures disease-associated changes in tTE and enables systematic characterization of translational adaptation across malignant and normal hematopoietic cell states.

### Life-span single-cell brain atlas reinforces neuron-specific tRNA signatures

Recent advances in single-cell technologies have enabled unprecedented resolution in the characterization of brain cell population and their molecular signatures (26–28). These efforts have accelerated the understanding of aging-associated processes and their contribution to neurodegenerative conditions, such as Alzheimer’s disease (AD) (29–31). To investigate translational regulation across neuronal and non-neuronal populations, we applied tTEscanR to scRNA-seq and scATAC-seq of mouse brains across lifespan (young, adult and aged) and two AD models (5xFAD and APOE) profiled in (30) **(Figure 3A; Supp. Figure 3A)**.

**Figure 3.**
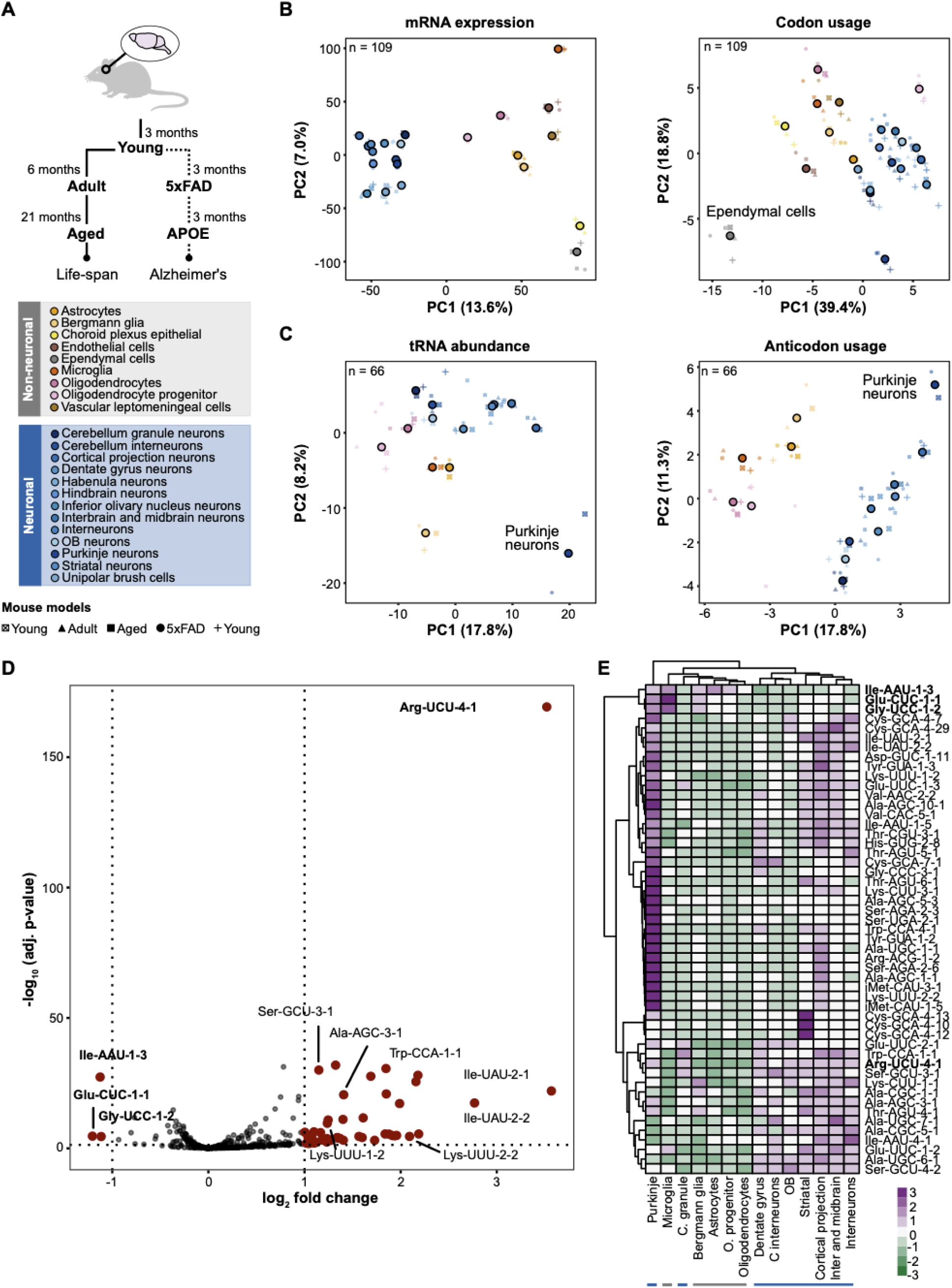
tTEscanR characterizes translational supply dynamics across lifespan and in Alzheimer’s disease models. **A.** Overview figure illustrates the five mouse models evaluated in this study, encompassing paired scRNA-seq and scATAC-seq data across three natural lifespan stages (young, adult, and aged) and two Alzheimer’s disease models (5xFAD, early onset: APOE, late onset). The pseudobulk strategy implemented defined conditions as the combination of mouse model and cell types. **B-C.** PCA plots visualize (**B**) mRNA gene expression (left) and codon usage (right), and (**C**) tRNA gene abundance (left) and anticodon usage (right). Data points are colored by cell type, with distinct shapes representing the different mouse models. Variations of blue denote specific neuron subtypes while other colors designate non-neuronal populations. Each point represents a pseudobulk condition defined by the mouse model and cell type combination with larger circles indicating median values. Clear separation between neuronal and non-neuronal lineages is evident in mRNA expression, tRNA abundance, and anticodon usage, but absent in codon usage, where ependymal cells uniquely deviate from the remaining population. **D.** Volcano plot compares neuronal and non-neuronal cell types across all mouse models. Statistical significance thresholds were set at | log_2_ | FC > 1 and -log_10_ adjusted *p*-value > 1.25. The analysis identified 51 tRNA genes with significantly altered tRNA gene usage in neurons with 48 enriched (positive log_2_ FC) and three (*Ile-AAU-1-3, Glu-CUC-1-1 and Gly-UCC-1-2*) depleted (negative log_2_ FC). **E.** Heatmap displays the expression patterns of the 51 deregulated tRNA genes identified in panel D across neuronal and non-neuronal cell types from all evaluated mouse models. tTEscanR resolved subtype-specific tRNA signatures within neuronal populations while preserving a shared pan-neuronal signature. The color scale represents row-normalized Z-scores (relative variance from the mean), with green indicating relative downregulation and purple indicating relative upregulation of tRNA abundance.

At the mRNA level, tTEscanR clearly separated neuronal cells from non-neuronal cell populations regardless of age or AD model status **(Figure 3B, left panel)**, reflecting the distinct transcriptional programs that define these cell types. In contrast, codon usage was overall more similar with neuronal and non-neuronal cell types aligning closer in the reduced-dimensional space **(Figure 3B, right panel; Supp. Figure 3B)**. Ependymal cells presented a notable deviation from this trend and displayed a modest divergence in codon usage relative to the remaining populations.

At the tRNA gene level, neuronal and non-neuronal cell populations were separated across all age groups and both AD models **(Figure 3C, left panel)**. Consistent with previous reports assessing cytosolic tRNA profiles (32), we did not observe differences in tRNA gene usage across aging or AD models, suggesting that cell identity is the dominant source of tRNA gene variation. Using the cell type annotations provided by the study, we found Purkinje neurons to cluster separately from the other cell types. At the anticodon level, separation of neuronal and non-neuronal cell populations became more pronounced **(Figure 3C, right panel; Supp. Figure 3C)**, consistent with previous observations in mouse and human brain tissues all neuronal subtypes clustered together and apart from non-neuronal cell types (19,33).

To further resolve these differences, we used tTEscanR to assess tRNA gene usage in neuronal and non-neuronal cell types. Because aging and AD models showed highly concordant patterns, we aggregated all datasets to maximize statistical power (**Methods**). This analysis resulted in 51 tRNA genes with significantly altered usage in neurons, including 48 enriched and three depleted tRNA genes (*Ile-AAU-1-3, Glu-CUC-1-1 and Gly-UCC-1-2*) **(Figure 3D; Supp. Figure 3D; Supp. Figure 4**). These neuron-associated tRNA gene signatures strongly overlapped with previously described neuronal tRNA programs identified in homeostatic brain tissues (19,33), supporting the robustness of tTEscanR across datasets and experimental systems (**Supp. Figure 3E**).

The high cellular resolution of this dataset further enabled tTEscanR to identify subtype-specific tRNA signatures within neuronal populations, while preserving a shared pan-neuronal signature **(Figure 3E)**. Most neuronal cell types shared a conserved tRNA profile, which was most pronounced in cortical projection neurons. However, cerebellum granule neurons exhibited a distinct pattern consistent with previous Pol III occupancy profiles (33). Notably, we observed that several neuronal populations showed selective enrichment of cysteine tRNA genes. For example, striatal and Purkinje neurons were enriched in six multiple tRNA^Cys^(GCA) genes (*Cys-GCA-4-7/10/12/13/29* and *Cys-GCA-7-1*). The strong enrichment likely reflects both the genomic clustering of cysteine tRNA genes and their coordinated regulation in neuronal translational programs.

Finally, we restricted our tTE analysis to the 66 matched conditions shared between the scRNA-seq and scATAC-seq datasets. Compared with our previous analyses (19), this dataset focused almost exclusively on brain cell types and therefore contained a relatively limited representation of non-neuronal cell types relative to the highly resolved neuronal subclasses. Besides this imbalance, tTEscanR detected a statistically significant enrichment of tTE scores at the codon-anticodon level in neuronal compared to non-neuronal cell types **(Supp. Figure 3F)**, further supporting the notion of enhanced translation efficiency in neurons.

## Discussion

In this study, we present tTEscanR, an open-source R package that enables scalable and reproducible analysis of codon and anticodon deployment for tTE from bulk and single-cell sequencing data. Building on our previously developed framework (19), tTEscanR extends the underlying methodology with a standardized software environment, integrated preprocessing and quality control, modular analytical components, and optimized computational performance while adhering to the FAIR principles (34). A central goal of our package development was to lower the technical barriers and automatize complex analytical procedures into an integrated workflow that requires minimal user input. To this end, tTEscanR streamlines complex computational procedures into a user-friendly workflow that requires only processed mRNA and tRNA count matrices as input, enabling researchers without extensive computational expertise to interrogate translational dynamics across biological models. At the same time, the modular design and open-source implementation allow advanced users to customize individual components and adapt the framework towards specialized applications.

Beyond the core algorithm, tTEscanR incorporates dedicated preprocessing, quality-control, and visualization modules that facilitate robust and reproducible analyses. Compared with our original scripts, extensive code optimization has been made to improve scalability for increasingly large single-cell atlases. Because the reliability of translation efficiency estimates ultimately depends on input data quality, we encourage the use of established preprocessing and quality-control guidelines adapted to each sequencing technology before applying tTEscanR. Appropriate filtering remains particularly important for sparse single-cell datasets, where technical noise can bias estimates of codon demand and anticodon supply and consequently affect downstream biological interpretation. To further address this challenge, tTEscanR implements adaptive thresholding procedures that move beyond fixed filtering criteria by identifying dataset-specific cutoffs that maximize retention of biologically informative conditions while maintaining stable anticodon quantification. This data-driven approach improves statistical power while minimizing unnecessary loss of information in heterogeneous datasets. Together, these features transform a previously custom analytical framework into a comprehensive software solution for studying codon demand, anticodon supply, and translation efficiency.

We have applied tTEscanR across both longitudinal patient cohort and large-scale tissue atlases to demonstrate its power to uncover coordinated translational programs at single-cell resolution. Across both human hematopoietic populations and the mouse brain, global codon usage (demand) remained remarkably stable across cell types and disease states. However, tTEscanR successfully captured distinct biological exceptions to this baseline. In the hematopoietic lineage, it revealed how the highly specialized transcriptomic program of erythroid cells, dominated by hemoglobin production, directly reshapes cell-specific codon demand. Conversely, within the central nervous system, codon demand remained uniform across neuronal subtypes. Nevertheless, tTEscanR robustly distinguished neuronal subtypes based on tRNA gene and anticodon accessibility (supply), recapitulating known cell type-specific tRNA signatures (19,33) and highlighting distinct tRNA profiles. Together, these lineage- and cell type-specific validations confirm the sensitivity of tTEscanR to map complex translational landscapes using standard multi-omic inputs.

Beyond mapping physiological cell states, tTEscanR exposed systematic shifts in the translational machinery during disease progression. In pAML, malignant myeloid cells displayed significantly elevated tTE scores compared to their remission counterparts, suggesting that leukemic populations optimize their codon-anticodon matching to support translational efficiency of proliferative or oncogenic networks. In contrast, aging and AD models exhibit unexpected stability in their tRNA supply landscapes. Rather than implying that translation is completely unaltered in neurodegeneration, where protein aggregation, cellular stress, and downstream degeneration are intricately linked (35), this stability indicates that disease-associated translational defects may be highly localized or subtle, remaining largely masked by the dominant molecular signatures of cell identity.

Several limitations should be considered. First, tTEscanR infers theoretical translation efficiency from codon demand and anticodon supply and therefore complements rather than replaces direct measurements of translation such as Ribo-seq. Second, accurate estimation of anticodon availability remains dependent on sequencing depth and data quality, particularly in sparse single-cell datasets. Although the preprocessing and quality control modules implemented in tTEscanR mitigate these challenges, careful interpretation remains necessary in low-coverage settings. Finally, additional layers of translational regulation, including tRNA modifications (13), aminoacylation status (36), ribosome heterogeneity (37–39), and codon context effects, are not explicitly modeled and represent important avenues for future development.

Overall, tTEscanR provides a robust and scalable computational platform to investigate the translational interface between the transcriptome and proteome. Because it operates directly from standard transcriptomic and chromatin accessibility count matrices, it can be readily incorporated into routine bulk and single-cell RNA-seq and ATAC-seq pipelines. Adding translation efficiency as a complementary analytical layer to conventional gene expression and chromatin accessibility analyses enables the identification of translational regulatory programs that would otherwise remain overlooked. Beyond the applications in cancer biology, aging, and neurodegeneration presented here, tTEscanR can support investigations in immunology, infectious diseases, developmental biology, regenerative medicine, and pharmacogenomics, as well as biotechnology, synthetic biology, and mRNA therapeutic design. Looking forward, integration of tTEscanR with large-scale perturbational approaches, including CRISPR screens and multi-omic profiling strategies, may uncover translational dependencies and vulnerabilities that contribute to cellular fitness, disease progression, and therapeutic response.

### Conclusions

In summary, tTEscanR provides a scalable and user-friendly R framework for analyzing codon usage, anticodon availability, and theoretical translation efficiency from bulk and single-cell sequencing data. Its modular design enables integration into standard transcriptomic and epigenomic workflows, making translation efficiency analysis accessible to both computational and non-computational users while supporting reproducible and customizable analyses.

By linking codon demand with tRNA-derived supply, tTEscanR extends conventional transcriptome analysis with a functional layer that captures translational regulation in health and disease. We anticipate that its routine use across high-throughput datasets will enable systematic discovery of context-specific translational programs and establish translation efficiency as a complementary dimension in genomic studies.

## Methods

### tTEscanR workflow overview

The tTEscanR package is a modular software platform that builds on and substantially expands the previously established theoretical translation efficiency (tTE) methodolgy (19). To ensure high software reliability and structural integrity, the package was developed in accordance with the architectural principles described before (40). A comprehensive list of core functions, including specific parameters and primary use cases, is provided in **Supp. Table 1**.

The tTEscanR architecture supports both a fully automated “one-step” execution and a stepwise workflow. While the wrapper function *runPipeline*() provides a unified entry point that automates the entire analytical trajectory, tTEscanR is intentionally decoupled into independent modules. This design allows researchers to execute specific segments of the pipeline based on their available data or specific biological questions.

To ensure high interoperability with the broader R bioinformatics ecosystem, tTEscanR supports standard classes, including *Seurat* (*41*) and *SummarizedExperiment*, allowing tTEscanR to function either as a standalone tool or as a plug-in component to existing analytical pipelines.

### tTEscanR object architecture

The core data structure of the package is the *tTEscanR_Object*, implemented as a formal S4 class to ensure rigorous data encapsulation. The object is primarily composed of two dedicated slots, *assays* and *meta.data*, both defined as lists to provide the flexibility required for heterogeneous multi-omic data. This architecture is designed to manage the high-dimensional requirements of both bulk and single-cell experiments while preventing data fragmentation. By utilizing an object-oriented framework, tTEscanR guarantees that all the data remains coupled with their corresponding metadata throughout the entire analytical workflow. The object is initialized via *createObject*() and dynamically maintained for continuous integration of new analytical layers through *updateObject*().

To facilitate automated processing and uphold that downstream functions can locate required data programmatically, tTEscanR enforces a strict naming convention for elements within its internal slots. Primary transcriptomics inputs must be stored under the reserved “mRNA” identifier within the assays list, while chromatin-derived tRNA data is stored as “tRNA”. As the analysis progresses, computed intermediate results are assigned specific labels. A comprehensive mapping of these reserved identifiers, including the specific functions responsible for their generation and their corresponding storage locations within the object, is provided in **Supp. Table 1**. This standardized labeling schema is automatically verified during object updates to maintain nomenclature consistency across the pipeline.

Primary assay inputs for the *tTEscanR_Object* consist of count matrices where genomic features are represented as rows and experimental conditions as columns. These features are categorized as protein-coding genes for mRNA assays and high-confidence predicted tRNA genes for tRNA assays. To ensure computational efficiency and accommodate the inherent memory requirements of large-scale datasets, the object is designed to support multiple data formats, including sparse matrices. Beyond the features, conditions are defined as sample-level labels used to identify each observation, defined as intersections of various metadata variables, such as tissue type, replicate, clinical stages, treatment group, and/or cell type.

### Dataset-specific tRNA cuts threshold

We developed a dedicated module to preprocess (sc)ATAC-seq data, facilitating the transition from a standard peak-level count matrix to the tRNA-specific expression matrix required by the tTEscanR object. Overall, this set of functions predict tRNA gene locations within the fragment coordinates provided in the peak matrix, facilitating accurate mapping, implementing the tRNA annotation guidelines originally described in (19). This process is executed in two stages. Initially, the *tRNAGetMatrix*() function maps chromatin fragments to genomic coordinates provided in a reference file of expected high confidence tRNA genes retrieved from GtRNAdb (5). Subsequently, the *tRNASetGenes*() function processes these overlaps to predict and annotate the tRNA genes.

A significant technical optimization in tTEscanR is the automated and dataset-specific determination of the tRNA cuts threshold via the *tRNASetCutoff*() function. This is particularly critical for sparse single-cell datasets where a universal cutoff, such as the 5,000-cut threshold suggested in (19) may be overly stringent, risking the loss of entire biological conditions. The selection algorithm operates through an iterative correlation analysis. First, the function calculates a reference baseline for anticodon usage and amino acid supply using the entire unfiltered matrix. For each feature ƒ, its reference baseline value (*R*_ƒ_) is calculated as its total count across all conditions divided by the global aggregate sum:

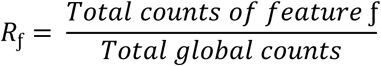

Next, the function tests a user-defined sequence of potential thresholds. For each tested value *t*, the data is subset to include only conditions that meet the minimum fragment count. The software then computes an “observed” profile for each iteration and measures its Spearman rank correlation coefficient (*ρ*) against the global reference baseline:

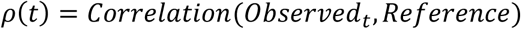

The final optimal threshold (*t* ∗) is determined by identifying the point of statistical stability. The algorithm evaluates the correlation coefficients alongside a slope-stability analysis to detect where the correlation scores begin to plateau. It computes the rate of change (m) in correlation between consecutive thresholds:

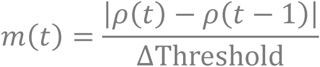

By selecting the final threshold at the onset of this stability, tTEscanR ensure that the data is sufficiently filtered to remove noise while retaining the maximum possible number of biological conditions for analysis.

### Baseline quantification of mRNA and tRNA data

The codon usage is inferred from transcriptome-wide expression data by parsing coding sequences into codon triplets and computing codon frequencies for each gene. These gene-level codon profiles are then weighted by transcript abundance (e.g., TPM or counts from RNA-seq or single-cell RNA-seq) and aggregated across all expressed genes to produce a sample-specific codon usage matrix that represents global translational demand (1,3,16). This weighting scheme ensures that highly expressed transcripts contribute proportionally more to the inferred codon landscape than lowly expressed genes.

On the anticodon side, tRNA-derived measurements are mapped to functional anticodon units through hierarchical aggregation of tRNA genes. Individual multi-copy tRNA genes are organized into isoacceptor families, defined by tRNA genes that share the same anticodon, and isotype classes, characterized by anticodons decoding the same amino acid. Analogous to codon usage estimation, tRNA-derived measurements are aggregated across tRNA genes sharing the same anticodon identity and weighted by their inferred activity, yielding sample-specific estimates of tRNA anticodon availability that enable systematic characterization of translational supply across diverse conditions and cell types.

### Automated generation of codon frequency tables

The quantification of codon usage is achieved through matrix multiplication between mRNA expression levels and a gene-specific codon frequency reference, as established in (19). This reference matrix captures the distribution of the 61 sense codons for every protein-coding gene within a specified genome. To enhance the versatility of tTEscanR, we implemented a module that allows users to generate these reference tables for any ds available via Ensembl or from custom FASTA sequences. This flexibility facilitates the application of the tTEscanR to non-model organisms or personalized genomic references beyond standard annotations. Furthermore, to fully implement the capabilities of a tailored codon usage analysis to all organisms we have included in tTEscanR the 33 available genetic codes available in NCBI Taxonomy (listed in **Supp. Table 1**).

The generation of the reference table is managed primarily through the *getCodonFreq*() function, which interfaces with the *biomaRt* package to retrieve genomic sequences (42). To maintain biological rigor, the function incorporates several filtering parameters to resolve transcript redundancy, allowing the user to isolate either the canonical transcript or the longest available sequence. Furthermore, multiple naming conventions are accommodated, including transcript identifiers, gene symbols, or Ensemble IDs. The software also permits the inclusion or exclusion of mitochondrial genes.

The *extractCodons*() function performs the raw counting of codon occurrences within a genomic sequence; while it remains available for direct execution by the user for custom sequences, it is primarily called internally by *getCodonFreq*() to format the final reference table. Because the extraction of genome-wide frequencies can be computationally intensive, tTEscanR provides pre-computed human (*hg38*) and mouse (*mm39*) reference tables as internal assets.

### Theoretical translation efficiency at the codon-anticodon level

The tTEscanR framework to compute the tTE scores builds upon the original tTE logic (19), which quantified translation efficiency based on the balance between amino acid supply and demand. In tTEscanR, we allow a higher resolution of tTE at the codon-anticodon level. However, due to the degeneracy of the genetic code, the 61 sense codons are recognized by a smaller pool of 46 anticodons. To evaluate this balance, the tTE score is calculated as the Spearman’s rank correlation coefficient between the supply vector (S) and demand vector (D) for each independent condition:

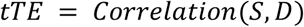

Because the framework relies on standard correlation metrics, the results tTE scores operate on a continuous scale bounded between −1 and 1:

*-1 ≤ tTE ≥ 1*

On this scale, a score approaching 1 indicates an optimal, highly coordinated balance between supply and demand, whereas a score near or below 0 reflects severe translational mismatching or disruption.

In this initial version of the tTEscanR algorithm, the mapping between these layers is restricted to canonical Watson-Crick complementary pairings. Consequently, non-canonical interactions such as wobble base-pairing rules or complex tRNA post-transcriptional modifications, are not currently incorporated into the scoring matrix.

### Additional modules within tTEscanR

To ensure seamless integration into existing bioinformatics ecosystems, the quantitative outputs generated by tTEscanR are structured as standardized high-dimensional matrices. This architecture is optimized not only for the package’s internal statistical modules, but also for direct export to external analytical pipelines in R and Bioconductor.

To facilitate the transition from raw quantification to statistical inference, the package includes dedicated functions for differential expression and permutation analysis and visualization options. By providing a suite of specialized plotting functions, tTEscanR enables users to summarize complex multi-omic findings in a visually accessible manner enhancing the overall clarity and impact of the analysis. A comprehensive list of visualization functions, including their specific parameters and primary use cases, is detailed in **Supp. Table 1**.

### Differential expression analysis

To identify shifts in the data, tTEscanR includes a differential expression module. This module is implemented through the *runDEAnalysis*() wrapper function, which sequentially executes *computeDEResults*() and *plotDEResults*(); notably, both functions remain available for independent execution to allow for modular workflows.

The *computeDEResults*() function integrates the DESeq2 framework (43). To account for technical variation, the function constructs a design formula based on user-defined condition and batch parameters extracted from the dataset metadata. This enables the model to systematically control for batch effects and other technical covariates while isolating condition-specific changes in the data. In the absence of known covariates, these parameters may be omitted to perform a univariate analysis.

tTEscanR facilitates two primary analytical modes: exploratory and targeted. The exploratory mode utilizes hierarchical clustering (heatmaps) and dimensionality reduction (PCA, UMAP, or tSNE) to identify global patterns and sample groupings. Conversely, the targeted mode allows for rigorous statistical comparisons between specific groups or a “one-vs-all” approach to identify cluster-specific signatures. These results are visualized via *plotDEResults*(). For targeted analysis, the function generates volcano plots to illustrate the magnitude (log_2_ fold-change) and statistical significance (-log_10_ adjusted *p*-value) of the findings. All visualization outputs are highly customizable (parameters detailed in **Supp. Table 1**).

### Permutation analysis

To evaluate whether specific functional gene sets exhibit specialized translational requirements, tTEscanR performs a permutation-based enrichment analysis. This module quantifies the divergence of the codon usage profile of a target gene set against the global transcriptomic baseline. Additionally, the module generates a density plot to visualize the observed enrichment relative to the null distribution.

The function *getPermutationDist*() generates a null distribution by randomly sampling gene sets of equivalent size *k* (where *k* equals the number of genes in the target set) from the background transcriptome. To ensure the independence of the background, any genes present in the target set are excluded from the sampling pool. For each iteration *i*, the raw codon frequencies are aggregated across the sampled subset and normalized to their relative contribution. For a given codon *c*, the normalized frequency *F_c_*_,*i*_ in permutation iteration *i* is calculated as:

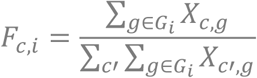

Where *G_i_* is the randomized subset of *k* background genes chosen during iteration *i*, and *X_c_*_,*g*_ represents the exonic count of codon *c* in gene *g*. By default, the algorithm performs *N* = 1,000 iterations to assemble the reference empirical null distribution **D*_c_*** = {*F_c_*_,1_, *F_c_*_,2_,…, *F_c_*_,*i*_}.

The function *obtainSignificance*() calculates empirical *p*-values by comparing the observed frequency (*f_obs_*) of each codon in the target dataset to its respective simulated null distribution **D***_c_*. To account for both enrichment and depletion, the function performs a conditional tail test relative to the median of the distribution:

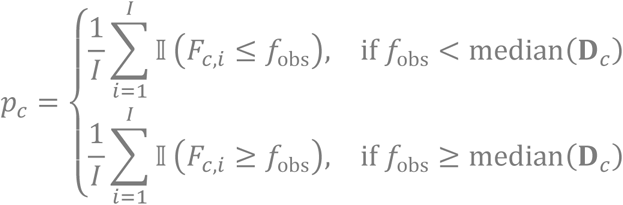

Where ‖()is an indicator function that equals 1 if the condition inside the parenthesis is met and 0 otherwise, and *I* represents the total number of permutations executed.

To control the False Discovery Rate (FDR) across the 64 codons, *p*-values are adjusted using the Benjamini-Hochberg (BH) method. Codons are considered significant if the adjusted *p*-value falls below a user-defined threshold (default *p_adj_*. < 0.05).

### tTEscanR pipeline execution

The analyses included in this manuscript were performed using the tTEscanR core pipeline with default parameters, unless otherwise specified. We utilized precomputed canonical human (*hg38*) and mouse (*mm39*) reference codon frequency tables as the transcriptomics baseline for the pAML and mouse brain cohorts, respectively.

### Pediatric AML cohort characteristics

The initial pediatric acute myeloid leukemia (pAML) cohort (25) comprised 28 patients. From this group, we prioritized a subset of 13 individuals (details provided in **Supp. Table 2**) with matched scRNA-seq and scATAC-seq data across three clinical stages: diagnosis, remission and relapse. Applying the unified preprocessing strategy detailed below, two patients were excluded from downstream analysis: AML11 failed to meet the minimum cell count threshold at the relapse stage, while AML21 exhibited a significant malignant burden during remission, preventing the establishment of reliable healthy baseline. Consequently, 11 patients were retained for the final tTEscanR analysis.

### Single-cell mouse brain atlas characteristics

The single-cell mouse brain atlas (30) spans five biological models with matched EasySci-RNA and EasySci-ATAC. This included a wild-type life-span cohort consisting of young (3 months), adult (6 months), and aged (21 months) mice. Additionally, the atlas included two Alzheimer’s disease (AD) phenotypes profiled at 3 months of age, representing models of early-onset (5xFAD) and late-onset (APOE) AD. Each of the five models was represented by four biological replicates, strictly balanced by sex (two males and two females) that were combined.

### Unified preprocessing strategy

Raw count matrices and associated metadata for both the pAML and mouse brain cohorts were kindly provided by the authors of the source studies (25,30). Gene expression matrices (scRNA-seq and EasySci-RNA) were processed following standard Seurat guidelines (41). For chromatin accessibility data (scATAC-seq and EasySci-ATAC), tRNA gene-per-cell count matrices were generated from the original peak matrices and fragment files using the dedicated tTEscanR module via the *tRNAGetMatrix*() *and tRNASetGenes*() functions. A quality control threshold of 2,724 (pAML) and 2,165 (mouse brain) tRNA cuts was determined using *tRNASetCutoff*() across 10,000 iterations and subsequently applied using *tRNAFilterCuts*().

### Dataset-specific preprocessing refinement

Following initial quality control, the datasets were refined based on the original study metadata.

#### pAML cohort

To facilitate a robust comparison between malignant and healthy states, we retained only malignant cells from the diagnosis and relapse stage, while restricting remission samples to healthy cell populations. Cells with unknown annotation were excluded.

#### Mouse brain cohort

To maximize read depth and statistical power, we utilized the “general” annotation provided in the metadata for initial count matrix labeling. Consistent with the original study’s findings, preliminary analysis revealed no significant sex-specific variance (data not shown). Consequently, data from both sexes were pooled using *mergeMatrices*(). To resolve specific neuronal signatures, we performed manual cluster aggregation using *groupConditions*() for certain populations (e.g. merging OB neurons 1, 2, and 3 into a single “OB neurons” category). The cell type clustering strategy followed is detailed in **Supp. Table 3**.

### Matrix consolidation and conditions definition

After dataset-specific filtering, individual count matrices were consolidated using *mergeMatrices()* to generate unified objects for each modality. Cells were then aggregated into distinct analytical conditions using *groupConditions*(). For the pAML cohort, conditions were defined by the intersection of clinical stages and cell type designations (e.g. diagnosis_basophil). For the mouse brain dataset, conditions were defined by the intersection of experimental models and cell type designations (e.g. young_interneuron).

To ensure statistical reliability, any cell cluster containing fewer than 100 individual cells was removed. Additionally in the mouse brain dataset the pituitary cells were removed as they were only represented in one of the models after filtration. These criteria yielded 54 (scRNA-seq) and 42 (scATAC-seq) conditions for the pAML dataset across 3 models, and 109 (EasySci-RNA) and 66 (EasySci-ATAC) conditions for the brain dataset across 5 models (detailed in **Supp. Table 2 and Supp. Table 3**).

### Analysis of codon pool diversity in erythrocytes

To evaluate the specialization of the codon landscape, we utilized the *showPoolContribution*() function to identify gene sets exerting the most significant influence on global codon usage profiles. For each condition, the algorithm correlates the specific frequencies of each codon against the global mean usage to determine the degree of divergence from the average transcriptomic baseline. The codon pool contribution is defined as the product of the normalized mRNA expression levels and the total codon count for each gene. For each condition, the top 10 genes with the highest codon pool contribution were extracted and correlated against the overall codon-to-mean usage values. By mapping these top-contributing genes, tTEscanR facilitates the detection of master transcripts that skew the global codon landscape toward specific translational requirements To identify lineage-specific codon usage profiles, we analyzed the remission stage taking early and late erythrocytes as the reference populations. We quantified the relative enrichment of each codon by calculating the log_2_ fold-change (FC) of the reference erythroid population against a lineage background, defined as the mean frequency of each codon across all non-erythroid cell types. The log_2_ FC was computed using a pseudocount (1 x 10^-6^) to ensure numerical stability:

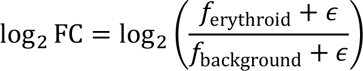

where *f* represents the codon frequency and *ε* is the pseudocount. A positive value indicates preferential enrichment in the erythroid lineage, while a negative value indicates relative under-representation.

To validate the influence of the erythroid program on global shifts, we profiled the ten hemoglobin complex genes present in the dataset (*HBA1*, *HBA2*, *HBB*, *HBD*, HBE1, *HBG1*, *HBG2*, *HBM*, *HBQ1* and *HBZ*). Coding sequences for these genes were retrieved from NCBI (accession numbers listed in **Supp. Table 2**), and their specific codon compositions were quantified using *extractCodons*(). We calculated the median codon counts across these reference genes to facilitate a direct comparison between hemoglobin-specific codon biases and the lineage-wide deviations observed in erythrocytes.

### tRNA signature profiling and cross-model normalization

We run a targeted differential expression analysis at the tRNA level comparing together all the models to interrogate neuronal (n = 9) against non-neuronal (n = 5) cell types using *runDEAnalysis*() with default parameters and correcting for the model and the targeted group (neurons vs non-neurons).

To characterize the expression patterns of the previously identified neuron-enriched tRNA genes (n = 51; absolute log_2_ FC > 1 and adjusted *p*-value < 0.05), we generated a consensus expression profile using a two-stage hierarchical averaging approach. First, to prevent biological bias from any individual model, data were first aggregated by calculating the mean expression of each tRNA within each model and cell type combination. Subsequently, a final consensus profile was then generated by averaging these model-specific means across all five biological groups (young, adult, aged, 5xFAD and APOE), resulting in a single representative value for each cell type. This normalization strategy ensures that the reported tRNA signatures are robust across the mouse lifespan and AD pathologies. For visualization, we generated a heatmap using the pheatmap package in R. To highlight the relative expression differences across cell types, data were scaled (Z-scores). Hierarchical clustering was preformed using correlation distance for the tRNAs and Euclidean distance for the cell types.

## Declarations

### Ethics approval and consent to participate

Not applicable, as all data used in this study were obtained from publicly available sources.

### Consent for publication

Not applicable.

### Availability of data and materials

All dataset analyzed in this study were retrieved from publicly available repositories. The raw sequencing data was downloaded from the Gene Expression Omnibus (GEO) under accession numbers GSE235063 (scRNA-seq), GSE235308 (scATAC-seq), GSM6538356 (EasySci-RNA) and GSM6538357 (EasySci-ATAC). The specific samples, clinical metadata, and cell state annotation evaluated in this study are detailed in **Supp. Table 2 and Supp. Table 3**.

The tTEscanR package is open-source and available from two sources: the primary development repository is **avarassanchez/tTEscanR** and the submission tracking for the Bioconductor review **Bioconductor/BiocContributions/issues/75**. The specific version used to generate the results in this manuscript is **v.0.99.0.** To ensure reproducibility, the repository includes comprehensive vignettes outlining the core module functionality, alongside all scripts and instructions required to preprocess the cohorts and replicate the figures presented herein.

## Acknowledgements

We are grateful to the BC CANCER consortia, Ziyu Lu and Junyue Cao for sharing invaluable data to the research community. This manuscript was prepared using a limited access dataset obtained from BC CANCER and does not necessarily reflect the opinions or views from BC CANCER. We sincerely appreciate the time and effort of Dr. Hugo Gruson in assessing our R package on behalf of Bioconductor.

We gratefully acknowledge the National Bioinformatics Infrastructure of Sweden (NBIS) Advisory Program for their constructive feedback on data analysis and data presentation. Part of the computations and data storage were enabled by resources provided by the National Academic Infrastructure for Supercomputing in Sweden (NAISS).

We thank the group members of the laboratories of Claudia Kutter, Vicent Pelechano and Marc Friedländer for their insightful feedback and friendly review.

---

**Supplementary Figure 1.**
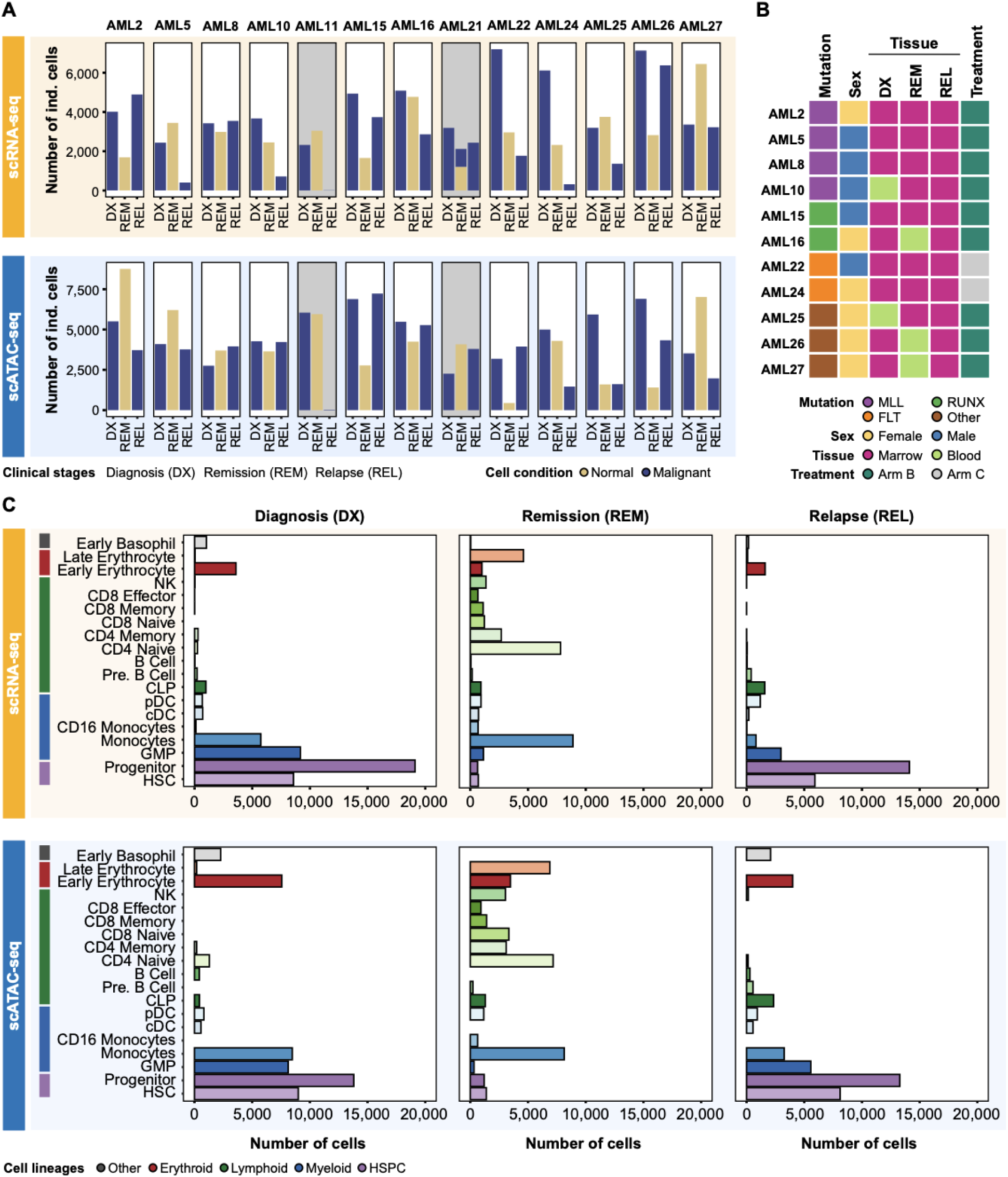
tTEscanR-based characterization of pAML cohort considers metadata information and cell count profiles. **A.** Barplots show the 13 patients (indicated by patient number) evaluated in this study containing paired scRNA-seq (top panel) and scATAC-seq (bottom panel) data across three clinical stages: diagnosis (DX), remission (REM), and relapse (REL). Diagnosis and relapse samples were filtered to retain only malignant cells (blue), whereas remission samples contain exclusively normal cells (gold). Patients highlighted in grey (AML11 and AML21) were excluded from downstream analyses due to failure in meeting quality filtering criteria. **B.** Metadata heatmap presents the 11 patients remaining after quality control and preprocessing, reflecting clinical and biological variations across mutational background, sex, tissue of origin of the sample, and treatment status. **C.** Horizontal barplots illustrate individual cell counts within major cell lineages (color-coded) retrieved from scRNA-seq (upper panel) and scATAC-seq (lower panel) datasets. Single cells were aggregated by cell type to implement a pseudobulk profiling strategy, and conditions for downstream analyses were defined by the combination of clinical stage and cell type of origin. The scATAC-seq counts correspond to the filtered dataset after applying the data-driven tTEscanR threshold of 2,724 tRNA gene-associated tags.

**Supplementary Figure 2.**
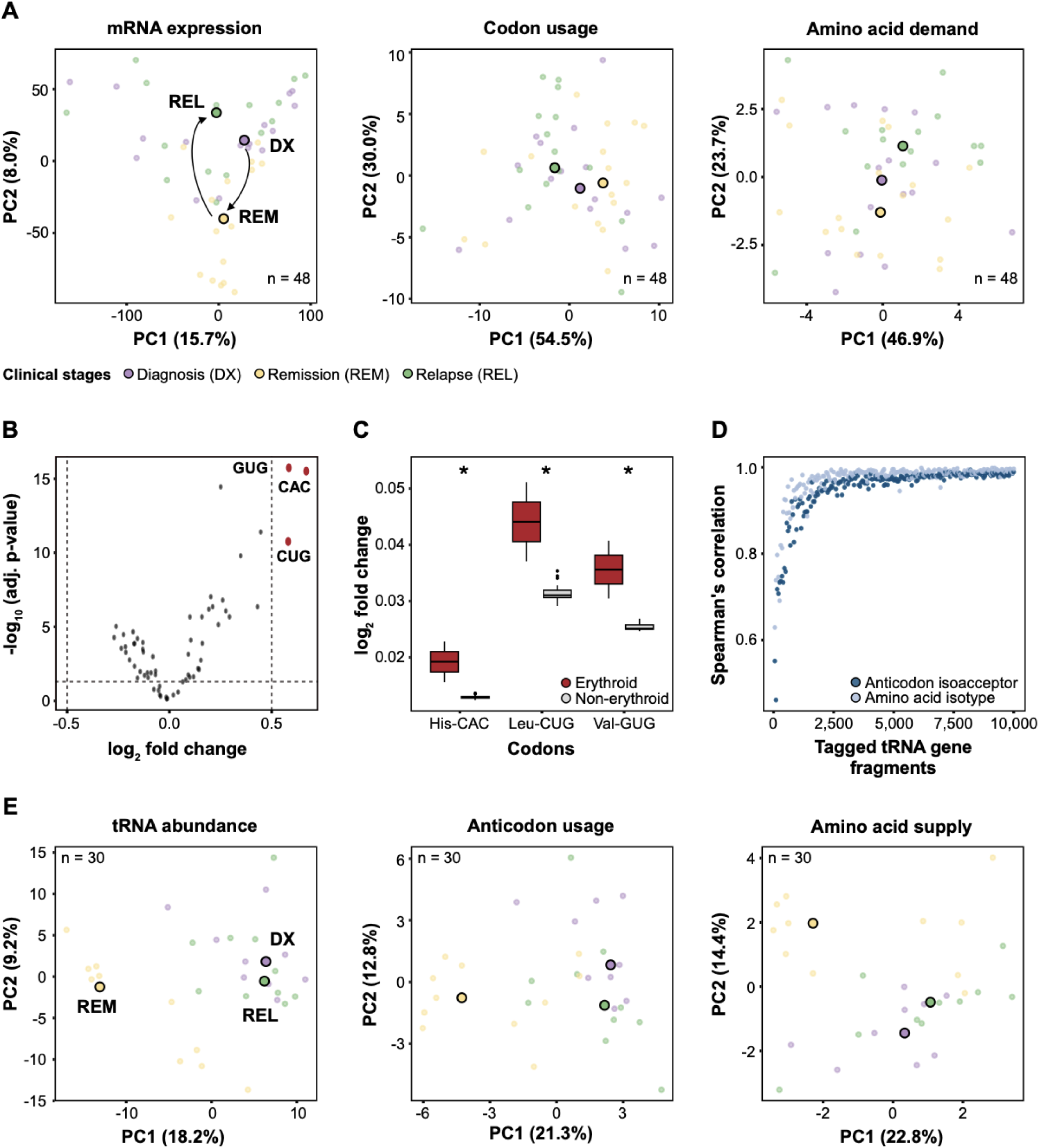
tTEscanR robustness testing validates codon enrichment in erythroid lineage and validates threshold optimization. **A.** PCA plots visualize mRNA gene expression (left), codon usage (center) and amino acid demand (right) after the removal of early and late erythrocytes, colored by clinical stage. Each data point represents a pseudobulk condition defined by the combination of clinical stage and cell types (n = 48, after filtering). Median values are highlighted with larger circles. **B.** Volcano plot illustrates codon usage in the erythroid lineage (early and late erythrocytes) against all the other cell types across the three clinical stages. Positive log_2_ fold-change (FC) values indicate enrichment in the erythroid lineage. The three most statistically significantly (log_2_ FC > 0.5) enriched codons are highlighted in red: Histidine-CAC, Leucine-CUG, and Valine-GUG. **C.** Box plots compare the relative abundance of the enriched codons identified in panel B between erythroid (red) and non-erythroid (grey) cell populations. Asterisks indicate statistical significance (*, *p*-value < 0.05). **D.** Correlation plot defines the optimal dataset-specific tTEscanR threshold by balancing signal robustness and data retention at the anticodon (dark blue) and amino acid supply (light blue) levels. In the pAML dataset, thresholds were tested over 10,000 iterations. The final optimal threshold of 2,724 tags was selected based on the point of maximal statistical stability. **E.** PCA plots visualize tRNA gene abundance (left), anticodon usage (center) and amino acid supply (right), colored by clinical stage. Data were filtered using the generic threshold of 5,000 tRNA gene-associated tags yielding a total of 30 conditions. Consistent with the tTEscanR-determined threshold (Fig. 2G), malignant cells from the diagnosis and relapse stages separate from normal hematopoietic populations in remission (yellow).

**Supplementary Figure 3.**
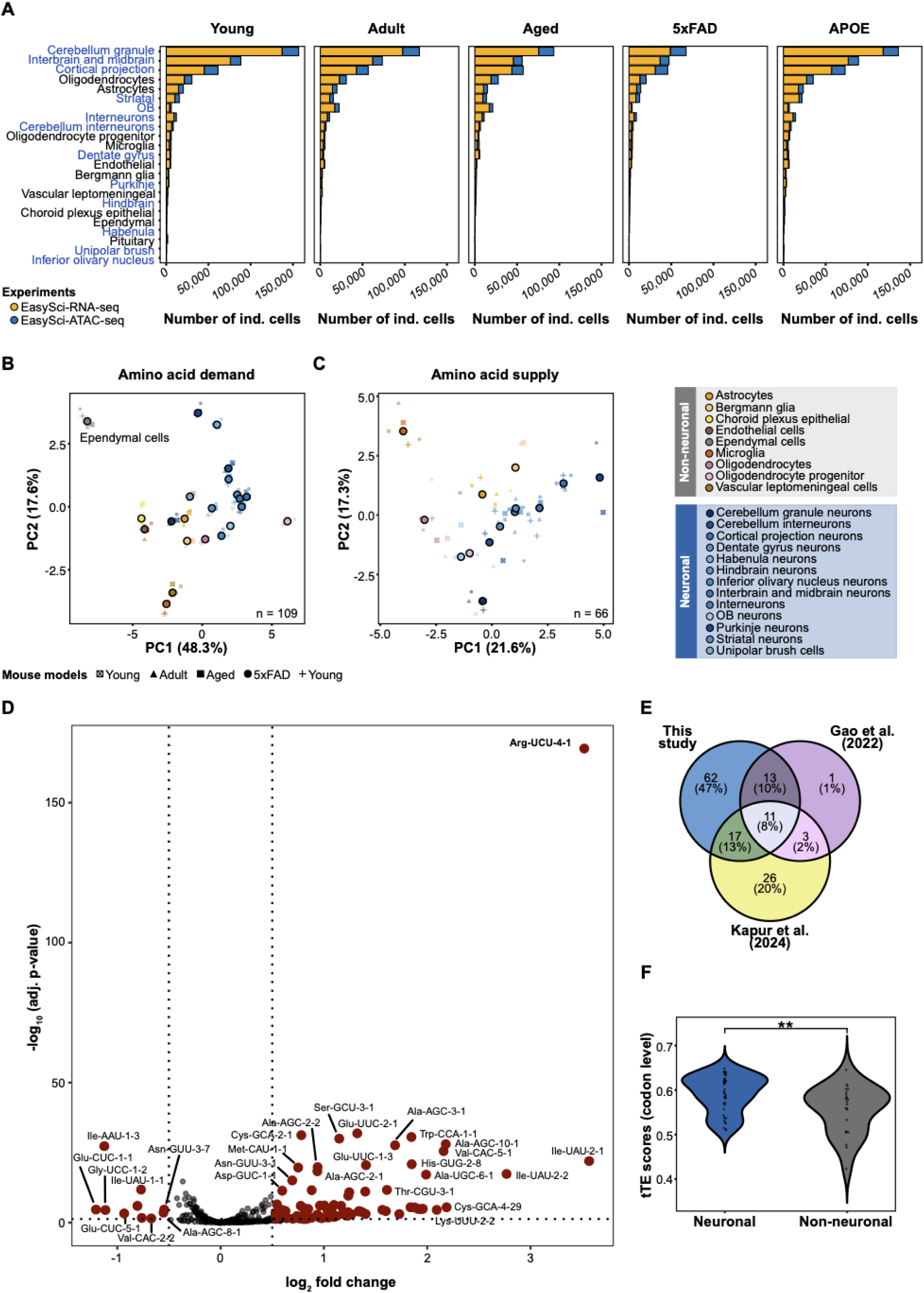
tTEscanR integrates pseudobulk profiling to resolve translational supply and demand across lifespan and disease models. **A.** Horizontal barplots illustrate individual cell counts used in the pseudobulk strategy to define analytical conditions based on the combination of mouse model and cell type of origin. scRNA-seq counts are represented in yellow, and scATAC-seq counts are shown in blue. Cell type labels on the y-axis are color-coded to distinguish neuronal populations (blue) from non-neuronal populations (black). **B-C.** PCA plots visualize amino acid **(B)** demand and **(C)** supply. Data points are colored by cell type, with distinct shapes representing the different mouse models. Variations of blue denote specific neuron subtypes while other colors designate non-neuronal populations. Each point represents a pseudobulk condition defined by a mouse model and cell type combination, with larger circles indicating median values. **D.** Volcano plot compares neuronal against non-neuronal cell types across all mouse models. Statistical significance thresholds were set at | log_2_ | FC > 0.5 and -log_10_ adjusted *p*-value > 1.25. The analysis identified 103 tRNA genes with significantly altered tRNA gene usage in neurons with 93 enriched (positive log_2_ FC) and 10 depleted (negative log_2_ FC). **E.** Venn diagram compares significantly altered tRNAs between neuronal and non-neuronal cell types from three sources: this study’s analysis of the mouse brain atlas (103 tRNAs), reanalysis of the adult mouse single-cell atlas (28 tRNAs) using tTEscanR: and a cell-type-specific Pol III epitope-tagging dataset (57 tRNAs). In total, 11 tRNA genes overlap across all three independent studies. **F.** Violin plot compares theoretical translation efficiency (tTE) scores between neuronal (blue) and non-neuronal (grey) cell types across all mouse models. Asterisks indicate statistical significance (**, *p*-value < 0.01) derived from a two-tailed standard normal distribution (Z-test).

**Supplementary Figure 4.**
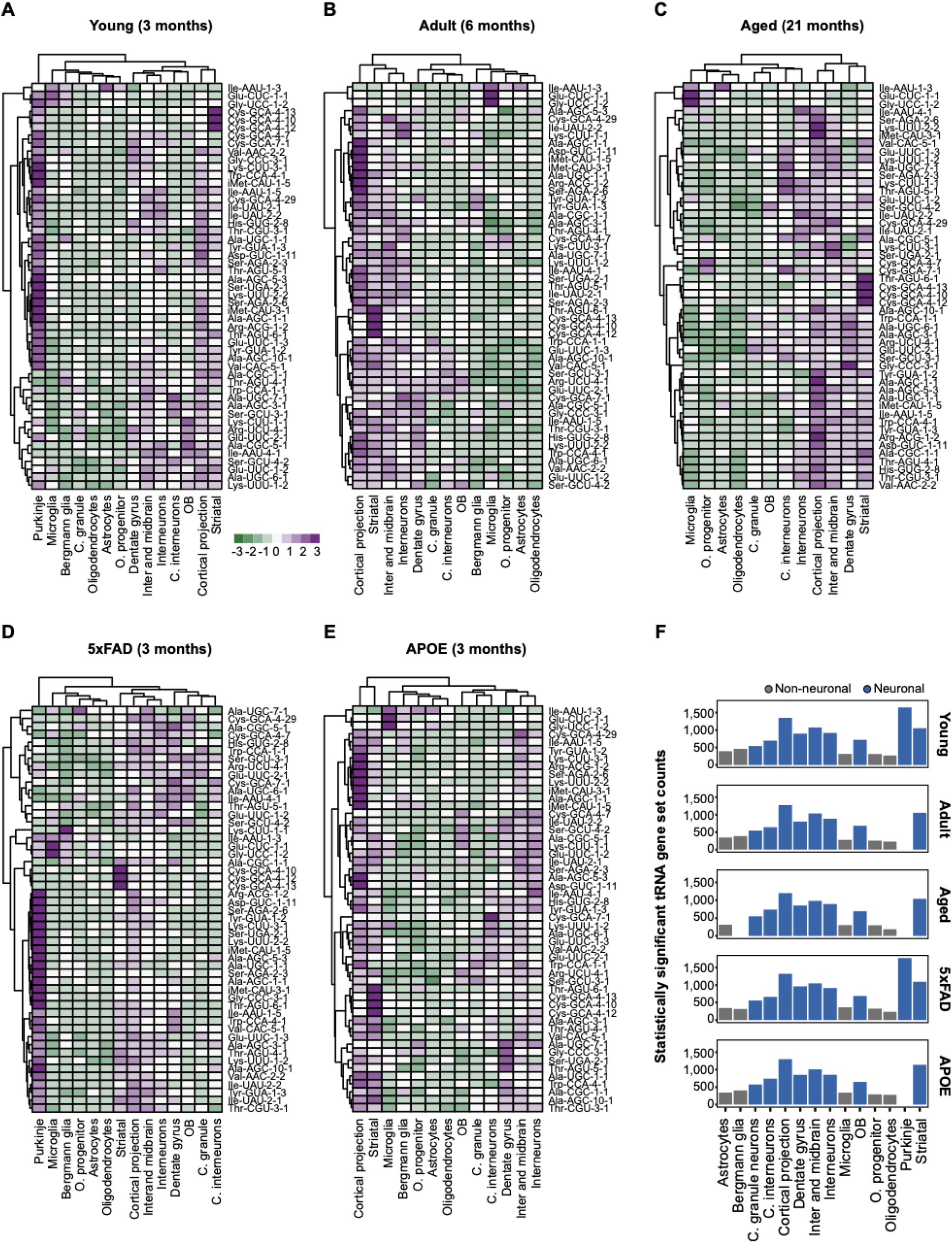
tTEscanR resolves stratified expression patterns and abundance profiles of deregulated neuronal tRNA genes. **A-E.** Heatmaps display the expression patterns of the 51 deregulated tRNA genes (| log_2_ | FC > 1 and -log_10_ adjusted *p*-value > 1.25) in neuronal versus non-neuronal cell lineages across individual mouse models: **(A)** young, **(B)** adult, **(C)** aged, **(D)** 5xFAD, and **(E)** APOE. To emphasize relative expression variations across cell lineages, data were row-normalized to calculate Z-scores, with green indicating relative downregulation and purple indicating relative upregulation of tRNA abundance. Hierarchical clustering was preformed using correlation distance for tRNAs and Euclidean distance for cell types. **F.** Barplots show normalized expression counts of the 48 upregulated tRNA genes in neurons across the evaluated models, demonstrating their elevated baseline abundance within specific cell types (neuronal cell types colored in blue; non-neuronal cell types in grey).

## References

1. Schmitt BM, Rudolph KLM, Karagianni P, Fonseca NA, White RJ, Talianidis I, et al. High-resolution mapping of transcriptional dynamics across tissue development reveals a stable mRNA–tRNA interface. Genome Res. 2014 Nov;24(11):1797–807. doi:10.1101/gr.176784.114

2. Saint-Léger A, Ribas De Pouplana L. The importance of codon–anticodon interactions in translation elongation. Biochimie. 2015 Jul;114:72–9. doi:10.1016/j.biochi.2015.04.013

3. Rudolph KLM, Schmitt BM, Villar D, White RJ, Marioni JC, Kutter C, et al. Codon-Driven Translational Efficiency Is Stable across Diverse Mammalian Cell States. PLOS Genet. 2016 May 11;12(5):e1006024. doi:10.1371/journal.pgen.1006024

4. Gallardo-Dodd CJ, Kutter C. The regulatory landscape of interacting RNA and protein pools in cellular homeostasis and cancer. Hum Genomics. 2024 Sep 27;18(1):109. doi:10.1186/s40246-024-00678-6

5. Chan PP, Lowe TM. GtRNAdb 2.0: an expanded database of transfer RNA genes identified in complete and draft genomes. Nucleic Acids Res. 2016 Jan 4;44(D1):D184–9. doi:10.1093/nar/gkv1309

6. Dittmar KA, Goodenbour JM, Pan T. Tissue-Specific Differences in Human Transfer RNA Expression. PLOS Genet. 2006 Dec 22;2(12):e221. doi:10.1371/journal.pgen.0020221

7. Pinkard O, McFarland S, Sweet T, Coller J. Quantitative tRNA-sequencing uncovers metazoan tissue-specific tRNA regulation. Nat Commun. 2020 Aug 14;11(1):4104. doi:10.1038/s41467-020-17879-x

8. Gao L, Behrens A, Rodschinka G, Forcelloni S, Wani S, Strasser K, et al. Selective gene expression maintains human tRNA anticodon pools during differentiation. Nat Cell Biol. 2024 Jan;26(1):100–12. doi:10.1038/s41556-023-01317-3

9. Schimmel P. The emerging complexity of the tRNA world: mammalian tRNAs beyond protein synthesis. Nat Rev Mol Cell Biol. 2018 Jan;19(1):45–58. doi:10.1038/nrm.2017.77

10. Gogakos T, Brown M, Garzia A, Meyer C, Hafner M, Tuschl T. Characterizing Expression and Processing of Precursor and Mature Human tRNAs by Hydro-tRNAseq and PAR-CLIP. Cell Rep. 2017 Aug 8;20(6):1463–75. doi:10.1016/j.celrep.2017.07.029

11. Shigematsu M, Honda S, Loher P, Telonis AG, Rigoutsos I, Kirino Y. YAMAT-seq: an efficient method for high-throughput sequencing of mature transfer RNAs. Nucleic Acids Res. 2017 May 19;45(9):e70. doi:10.1093/nar/gkx005

12. Behrens A, Rodschinka G, Nedialkova DD. High-resolution quantitative profiling of tRNA abundance and modification status in eukaryotes by mim-tRNAseq. Mol Cell. 2021 Apr;81(8):1802–1815.e7. doi:10.1016/j.molcel.2021.01.028

13. Suzuki T. The expanding world of tRNA modifications and their disease relevance. Nat Rev Mol Cell Biol. 2021 Jun;22(6):375–92. doi:10.1038/s41580-021-00342-0

14. Lucas MC, Pryszcz LP, Medina R, Milenkovic I, Camacho N, Marchand V, et al. Quantitative analysis of tRNA abundance and modifications by nanopore RNA sequencing. Nat Biotechnol. 2024 Jan;42(1):72–86. doi:10.1038/s41587-023-01743-6

15. Padhiar NH, Katneni U, Komar AA, Motorin Y, Kimchi-Sarfaty C. Advances in methods for tRNA sequencing and quantification. Trends Genet. 2024 Mar 1;40(3):276–90. doi:10.1016/j.tig.2023.11.001

16. Kutter C, Brown GD, Gonçalves Â, Wilson MD, Watt S, Brazma A, et al. Pol III binding in six mammals shows conservation among amino acid isotypes despite divergence among tRNA genes. Nat Genet. 2011 Oct;43(10):948–55. doi:10.1038/ng.906

17. White RJ. Transcription by RNA polymerase III: more complex than we thought. Nat Rev Genet. 2011 Jul;12(7):459–63. doi:10.1038/nrg3001

18. Van Bortle K, Marciano DP, Liu Q, Chou T, Lipchik AM, Gollapudi S, et al. A cancer-associated RNA polymerase III identity drives robust transcription and expression of snaR-A noncoding RNA. Nat Commun. 2022 May 30;13(1):3007. doi:10.1038/s41467-022-30323-6

19. Gao W, Gallardo-Dodd CJ, Kutter C. Cell type–specific analysis by single-cell profiling identifies a stable mammalian tRNA–mRNA interface and increased translation efficiency in neurons. Genome Res. 2022 Jan 10;32(1):97–110. doi:10.1101/gr.275944.121 PubMed PMID: 34857654.

20. Dieci G, Fiorino G, Castelnuovo M, Teichmann M, Pagano A. The expanding RNA polymerase III transcriptome. Trends Genet. 2007 Dec 1;23(12):614–22. doi:10.1016/j.tig.2007.09.001

21. Ingolia NT, Ghaemmaghami S, Newman JRS, Weissman JS. Genome-Wide Analysis in Vivo of Translation with Nucleotide Resolution Using Ribosome Profiling. Science. 2009 Apr 10;324(5924):218–23. doi:10.1126/science.1168978

22. Brar GA, Weissman JS. Ribosome profiling reveals the what, when, where and how of protein synthesis. Nat Rev Mol Cell Biol. 2015 Nov;16(11):651–64. doi:10.1038/nrm4069

23. Wang Q, Mao Y. Principles, challenges, and advances in ribosome profiling: from bulk to low-input and single-cell analysis. Adv Biotechnol. 2023 Dec 1;1(4):6. doi:10.1007/s44307-023-00006-4

24. VanInsberghe M, Van Oudenaarden A. Sequencing technologies to measure translation in single cells. Nat Rev Mol Cell Biol. 2025 May;26(5):337–46. doi:10.1038/s41580-024-00822-z

25. Lambo S, Trinh DL, Ries RE, Jin D, Setiadi A, Ng M, et al. A longitudinal single-cell atlas of treatment response in pediatric AML. Cancer Cell. 2023 Dec 11;41(12):2117–2135.e12. doi:10.1016/j.ccell.2023.10.008 PubMed PMID: 37977148.

26. Braun E, Danan-Gotthold M, Borm LE, Lee KW, Vinsland E, Lönnerberg P, et al. Comprehensive cell atlas of the first-trimester developing human brain. Science. 2023 Oct 13;382(6667):eadf1226. doi:10.1126/science.adf1226

27. Lu Z, Zhang M, Lee J, Sziraki A, Anderson S, Zhang Z, et al. Tracking cell-type-specific temporal dynamics in human and mouse brains. Cell. 2023 Sep 28;186(20):4345–4364.e24. doi:10.1016/j.cell.2023.08.042 PubMed PMID: 37774676.

28. Siletti K, Hodge R, Mossi Albiach A, Lee KW, Ding SL, Hu L, et al. Transcriptomic diversity of cell types across the adult human brain. Science. 2023 Oct 13;382(6667):eadd7046. doi:10.1126/science.add7046

29. Mathys H, Davila-Velderrain J, Peng Z, Gao F, Mohammadi S, Young JZ, et al. Single-cell transcriptomic analysis of Alzheimer’s disease. Nature. 2019 Jun;570(7761):332–7. doi:10.1038/s41586-019-1195-2

30. Sziraki A, Lu Z, Lee J, Banyai G, Anderson S, Abdulraouf A, et al. A global view of aging and Alzheimer’s pathogenesis-associated cell population dynamics and molecular signatures in human and mouse brains. Nat Genet. 2023 Dec;55(12):2104–16. doi:10.1038/s41588-023-01572-y

31. Jin K, Yao Z, van Velthoven CTJ, Kaplan ES, Glattfelder K, Barlow ST, et al. Brain-wide cell-type-specific transcriptomic signatures of healthy ageing in mice. Nature. 2025 Feb;638(8049):182–96. doi:10.1038/s41586-024-08350-8

32. Kristen M, Lander M, Kilz LM, Gleue L, Jörg M, Bregeon D, et al. DORQ-seq: high-throughput quantification of femtomol tRNA pools by combination of cDNA hybridization and Deep sequencing. Nucleic Acids Res. 2024 Oct 14;52(18):e89. doi:10.1093/nar/gkae765

33. Kapur M, Molumby MJ, Guzman C, Heinz S, Ackerman SL. Cell-type-specific expression of tRNAs in the brain regulates cellular homeostasis. Neuron. 2024 May 1;112(9):1397–1415.e6. doi:10.1016/j.neuron.2024.01.028

34. Barker M, Chue Hong NP, Katz DS, Lamprecht AL, Martinez-Ortiz C, Psomopoulos F, et al. Introducing the FAIR Principles for research software. Sci Data. 2022 Oct 14;9(1):622. doi:10.1038/s41597-022-01710-x

35. Zhang J, Zhang Y, Wang J, Xia Y, Zhang J, Chen L. Recent advances in Alzheimer’s disease: mechanisms, clinical trials and new drug development strategies. Signal Transduct Target Ther. 2024 Aug 23;9(1):211. doi:10.1038/s41392-024-01911-3

36. Giegé R, Eriani G. The tRNA identity landscape for aminoacylation and beyond. Nucleic Acids Res. 2023 Feb 28;51(4):1528–70. doi:10.1093/nar/gkad007

37. Genuth NR, Barna M. The Discovery of Ribosome Heterogeneity and Its Implications for Gene Regulation and Organismal Life. Mol Cell. 2018 Aug;71(3):364–74. doi:10.1016/j.molcel.2018.07.018

38. Gay DM, Lund AH, Jansson MD. Translational control through ribosome heterogeneity and functional specialization. Trends Biochem Sci. 2022 Jan;47(1):66–81. doi:10.1016/j.tibs.2021.07.001

39. Alkan F, Wilkins OG, Hernández-Pérez S, Ramalho S, Silva J, Ule J, et al. Identifying ribosome heterogeneity using ribosome profiling. Nucleic Acids Res. 2022 Sep 9;50(16):e95–e95. doi:10.1093/nar/gkac484

40. Wickham H. R Packages. 2nd ed. Sebastopol: O’Reilly Media, Incorporated; 2023. 1 p.

41. Hao Y, Stuart T, Kowalski MH, Choudhary S, Hoffman P, Hartman A, et al. Dictionary learning for integrative, multimodal and scalable single-cell analysis. Nat Biotechnol. 2024 Feb;42(2):293–304. doi:10.1038/s41587-023-01767-y

42. Durinck S, Spellman PT, Birney E, Huber W. Mapping identifiers for the integration of genomic datasets with the R/Bioconductor package biomaRt. Nat Protoc. 2009 Aug;4(8):1184–91. doi:10.1038/nprot.2009.97

43. Love MI, Huber W, Anders S. Moderated estimation of fold change and dispersion for RNA-seq data with DESeq2. Genome Biol. 2014 Dec 5;15(12):550. doi:10.1186/s13059-014-0550-8

